# Biofilm Lithography: High-resolution cell patterning via optogenetic adhesin expression

**DOI:** 10.1101/226027

**Authors:** Xiaofan Jin, Ingmar H. Riedel-Kruse

## Abstract

Bacterial biofilms represent a promising opportunity for engineering of microbial communities. However our ability to control spatial structure in biofilms remains limited. Here we engineer *Escherichia coli* with a light-activated transcriptional promoter to optically regulate adhesin gene expression. When illuminated with patterned blue light, long-term viable biofilms with spatial resolution down to 25*μm* can be formed on a variety of substrates and inside enclosed culture chambers without the need for surface pretreatment. A biophysical model suggests the patterning mechanism involves stimulation of transiently surface-adsorbed cells, lending new evidence to a previously proposed role of adhesin expression during natural biofilm maturation. Overall, this tool – termed ‘Biofilm Lithography’ – has distinct advantages over existing cell-depositing and patterning methods and provides the ability to grow structured biofilms, with applications towards an improved understanding natural biofilm communities, as well as the engineering of living biomaterials and bottom-up approaches to microbial consortia design.

---

Biofilms are surface-attached communities of microbes, and represent the predominant mode of life for bacteria on earth(1). While well known for their role in biofouling and infections(2, 3), biofilms can also be harnessed as biotechnological tools, such as those used in wastewater treatment(4). Recent research has highlighted their potential as living biomaterials in applications including the prevention of biofouling(5), nanoparticle templating, protein immobilization, and bioelectricity(6, 7), as well as a promising platform upon which to engineer synthetic microbial communities(8, 9).

A key feature of natural biofilms biofilms is distinct spatial patterning coupled to ecological relationships within the microbial community, such as metabolic division of labour between co-localized strains(10). In some cases this type of structure allows simultaneous biochemical reactions to occur that would be incompatible within single cells(11). Clearly, the full biotechnological capabilities of engineered beneficial biofilms cannot be realized without reliable tools to control biofilm structure.

Such patterning tools should ideally be able to structure stable/viable biofilms with high spatial resolution, in a variety of environments without necessarily requiring substrate pretreatment or directly exposed surfaces. Many techniques exist to pattern cells to surfaces, including (but not limited to) inkjet printing(12, 13), microcontact printing(14, 15), microfluidics(16), PDMS stenciling(17), patterned substrate modification(18–20), photoactivated antibiotic(21), light-switchable adhesion proteins(22), and optogenetic cyclic-di-GMP regulation(23, 24). Optogenetic approaches have also been used to control gene expression on pre-existing bacterial lawns (25). However, to our knowledge, no existing method comprehensively fulfills the listed requirements for biofilm patterning.

Here we present a new technique termed ‘Biofilm Lithography’ to structure bacterial biofilms by projecting optical patterns which induce a planktonic-to-biofilm phenotypic switch in engineered bacteria. Our method rests on two key elements: (1) Light regulated transcriptional elements have been developed for bacteria, such as pDawn, which uses a light-oxygen-voltage domain to regulate gene expression according to blue light illumination(26) (2) The switch from planktonic to biofilm phenotype in bacteria can be controlled by the expression of membrane proteins that promote cell-substrate attachment, such as antigen 43 (Ag43), a homodimerizing autotransporter adhesin which has been demonstrated to induce both biofilm formation as well as cell-cell adhesion(27, 28). Here we bring Ag43 under the control of pDawn, allowing us to pattern biofilms with light.

## Results

### pDawn-Ag43 expression drives light-regulated biofilm formation in *E. coli*

Based on our current understanding of adhesin expression in biofilms and light activated transcriptional elements, we postulated that *E. coli* cells expressing Ag43 from the blue-light-responsive promoter pDawn should transition from a planktonic to biofilm phenotype when illuminated by blue light. To test our hypothesis, we designed a construct (termed pDawn-Ag43) where a ribosomal binding site and the Ag43 coding sequence have been inserted downstream of the pDawn transciptional control element (Fig. 1a), and we then transformed pDawn-Ag43 into the MG1655 strain of *E. coli*, a weak native biofilm former(29) that is known to form biofilm with Ag43 overexpression(28).

**Fig. 1.**
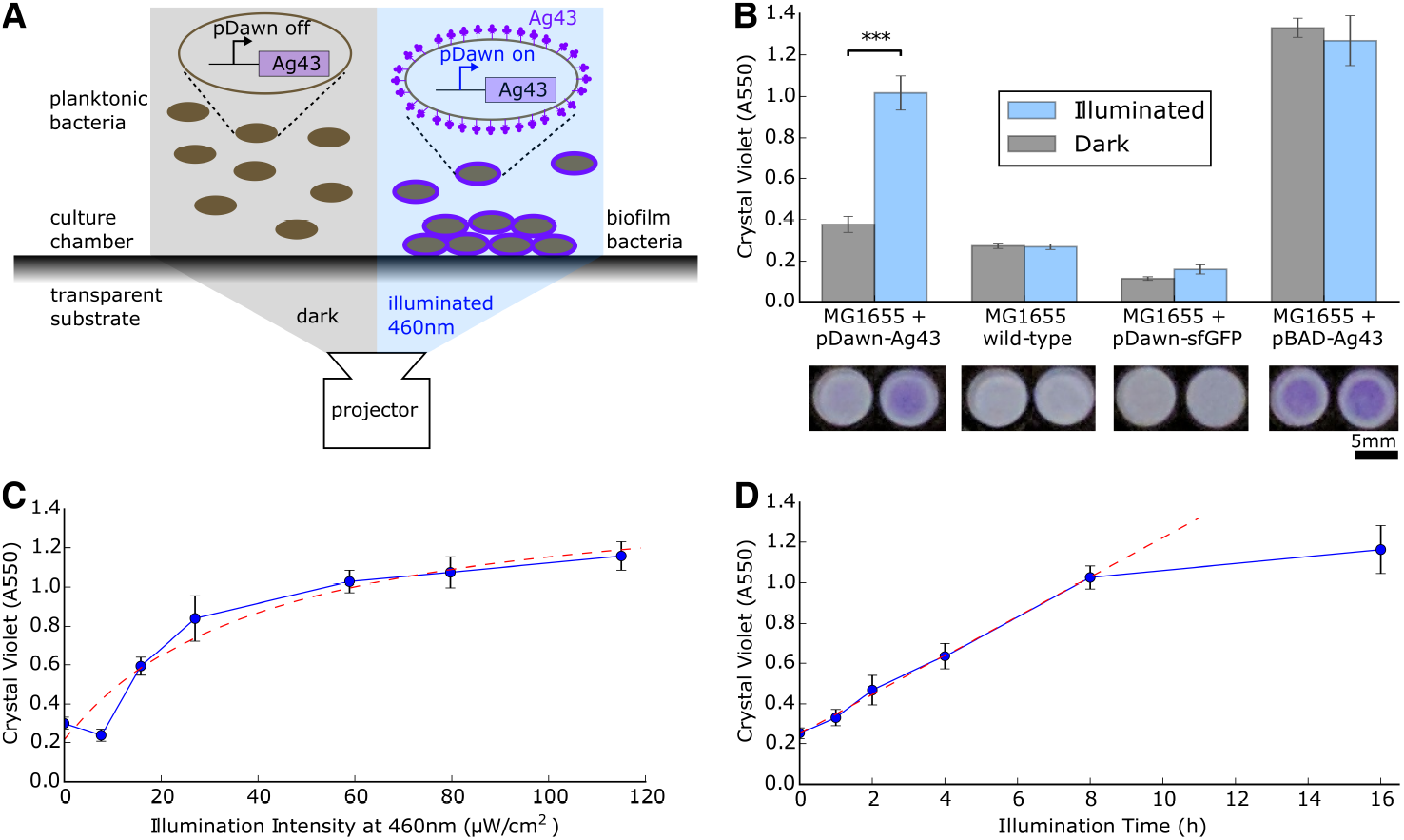
pDawn control of the adhesin Ag43 enables light-controlled deposition of biofilm onto surfaces. **(A)** Ag43 was inserted downstream of pDawn control. *E. coli* with pDawn-Ag43 can be optically stimulated with a projector in order to express Ag43 and to form biofilm at surfaces. **(B)** Cells engineered with the pDawn-Ag43 construct form biofilm contingent upon illumination. Wild-type and pDawn controls fail to form significant biofilm, and pBAD-Ag43 expression forms biofilm regardless of illumination. Representative 96-well crystal violet stains shown below. **(C)** Increasing illumination intensity increases biofilm formation, saturating past approximately 41 μW/cm^2^. Dashed curve: Fit with Monod model. **(D)** Increasing illumination time increases biofilm formation up to approximately 8h. Dashed curve: Linear model. (*B-D: STD error bars, n=4 wells*.)

We seeded cultures of MG1655/pDawn-Ag43 inside a polystyrene well-plate with M63 growth media, and used a portable LED projector (Ivation Pro4) to provide blue light illumination during an overnight growth incubation (Fig. 1a). Using crystal violet staining for quantification(30), we found that MG1655 transformed with pDawn-Ag43 formed robust biofilm when grown under illumination, compared to weak biofilm when grown in the dark (Fig. 1b).

To verify that the pDawn-regulated Ag43 expression is in fact responsible for this light dependent phenotype, we ran a series of controls: (1) untransformed MG1655, (2) MG1655 transformed with pDawn-sfGFP – a pDawn plasmid expressing superfolder green fluorescent protein (sfGFP) instead of Ag43, and (3) MG1655 transformed with pBAD-Ag43 – a plasmid expressing Ag43 under the control of the *p_BAD_* promoter, induced with 10mM arabinose (Fig. 1b). As expected, native MG1655 and MG1655 transformed with pDawn-sfGFP both failed to form strong biofilm regardless of dark or illuminated growth conditions. This indicates that at the intensity used (40*μ*W/cm^2^), blue light illumination on its own is unable to stimulate biofilm growth in MG1655, even with stimulation of the YF1/fixJ two component sensor involved in pDawn. We verified that pDawn works as expected in MG1655/pDawn-sfGFP by confirming sfGFP expression (Supplementary Information S1). On the other hand, MG1655 transformed with pBAD-Ag43 formed biofilm regardless of illumination intensity, indicating that Ag43 expression is sufficient to induce biofilm formation. We conclude that pDawn-regulated expression of Ag43 is responsible for the light-dependent biofilm phenotype in MG1655/pDawn-Ag43.

### Biofilm formation can be tuned by illumination intensity and time

We characterized the influence of illumination intensity and time on the extent of biofilm formation. By stimulating cells overnight for 16h across a range of intensities from 0 to 115*μ*W/cm^2^, we find that biofilm formation increases with brighter illumination in a saturating manner with a characteristic illumination intensity constant of 41*μ*W/cm^2^ when fit with the Monod equation (Fig. 1c, see fit/model choice discussion in Supplementary Information S2). When illuminating with 40*μ*W/cm^2^, we found a linear increasing relationship between biofilm formation and illumination time up to approximately 8h, beyond which biofilm formation appears to slow down (Fig. 1d). For the remainder of this paper we use standard illumination conditions of 40*μ*W/cm^2^ for 16 h (unless stated otherwise).

### Biofilm can be patterned using light in various environments

Next, we developed a protocol to form and visualize patterned biofilms based on pDawn-Ag43. We cotransformed MG1655/pDawn-Ag43 with a red fluorescent protein (mRFP) expression plasmid. Applying the same culture conditions as before, we set up the projector using Microsoft Powerpoint to illuminate various patterns (e.g., stripes, polka dots, pictures) on the bacterial samples, and subsequently verified that the patterns are recapitulated as bacterial biofilms using fluorescence microscopy (Fig. 2a).

**Fig. 2.**
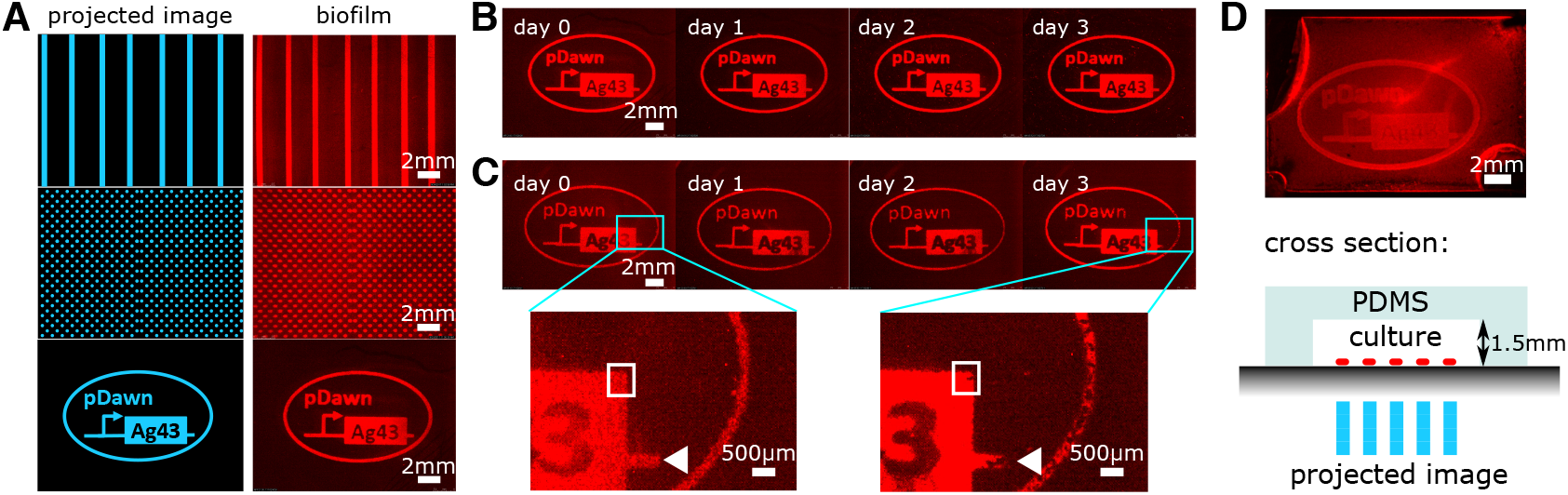
Biofilm Lithography enables the structured patterning of viable biofilms on various surfaces and inside closed culture chambers. **(A)** Various illumination patterns (stripes, polka dots, pictures) are recapitulated as biofilm patterns; projector patterns shown for reference. **(B)** Patterned biofilms remain stable for at least 3 days under ambient conditions in PBS buffer. **(C)** In culture media, growth and eventual detachment of viable biofilm can be observed over the same time span. White rectangles indicate regions with increased mRFP signal, triangles point to instances of biofilm detachment. **(D)** Biofilms can be patterned with light inside sealed, transparent culture chambers, without the need for direct physical access to the substrate. Within enclosed chamber, planktonic cells are more challenging to rinse without biofilm perturbation, resulting in reduced image contrast.

##### Significance Statement

Bacteria live in surface-attached communities known as biofilms, where spatial structure is tightly linked to community function. We have developed a genetically encoded biofilm patterning tool by engineering bacteria such that the expression of membrane adhesion proteins responsible for surface attachment is optically regulated. Accordingly, these bacteria only form biofilm on illuminated surface regions. With this tool, we are able to use blue light to pattern *Escherichia coli* biofilms with 25*μm* spatial resolution. We present an accompanying biophysical model to understand the mechanism behind light-regulated biofilm formation and to provide insight on related natural biofilm processes. Overall, this biofilm patterning tool can be applied to study natural microbial communities as well as to engineer living biomaterials.

We next investigated the long term stability of the patterned biofilms. After overnight incubation, samples were removed from optical illumination and rinsed with PBS, and subsequent daily imaging demonstrated that the biofilm pattern remained stable over a period of three days in PBS (Fig. 2b). When biofilm is maintained in M63 growth medium over the same time span, we observe growth and shedding of viable biofilm (Fig. 2c), analogous to expansion and dispersal processes in natural biofilms(29).

Next, we tested this tool’s ability to pattern other materials besides polystyrene. Using glass coverslips and PDMS coupons placed into well-plates, we confirm that our engineered cells are able to form patterned biofilms on both glass and PDMS (Supplementary Information S3). For all tested substrates, no surface pretreatment or patterning was required.

Furthermore, we tested whether our method works on samples inside completely enclosed environments. This is in contrast to the requirement of direct surface access for other patterning methods such as inkjet-based printers or microcontact printing (12–15). We used molded PDMS cavities bonded to polystyrene to create enclosed culture chambers with dimensions 19mm × 13mm × 1.5mm (Fig. 2d). Bacteria were cultured in this chamber using the same illumination conditions as before. We found that patterned biofilms form as expected inside enclosed environments (Fig. 2d).

### Biofilms can be optically patterned with 25*μm* resolution

In order to more quantitatively characterize pDawn-Ag43 mediated biofilm patterning, we collected volumetric biofilm data via confocal laser scanning microscopy (Fig. 3a), and measured the biofilm’s average thickness to be 14.4*μm*. (Supplementary Information S4). Assuming an average *E. coli* cell volume of 1*μm*^3^(31) and that cells constitute 10% of total biofilm volume(32), this approximately corresponds to an average thickness estimate of 7 cells and a surface density of 1.4x10^6^cells/mm^2^. From this we can estimate an average biofilm deposition rate on the order of 25 cells/mm^2^/s over the course of a 16h incubation (Supplementary Information S4).

**Fig. 3.**
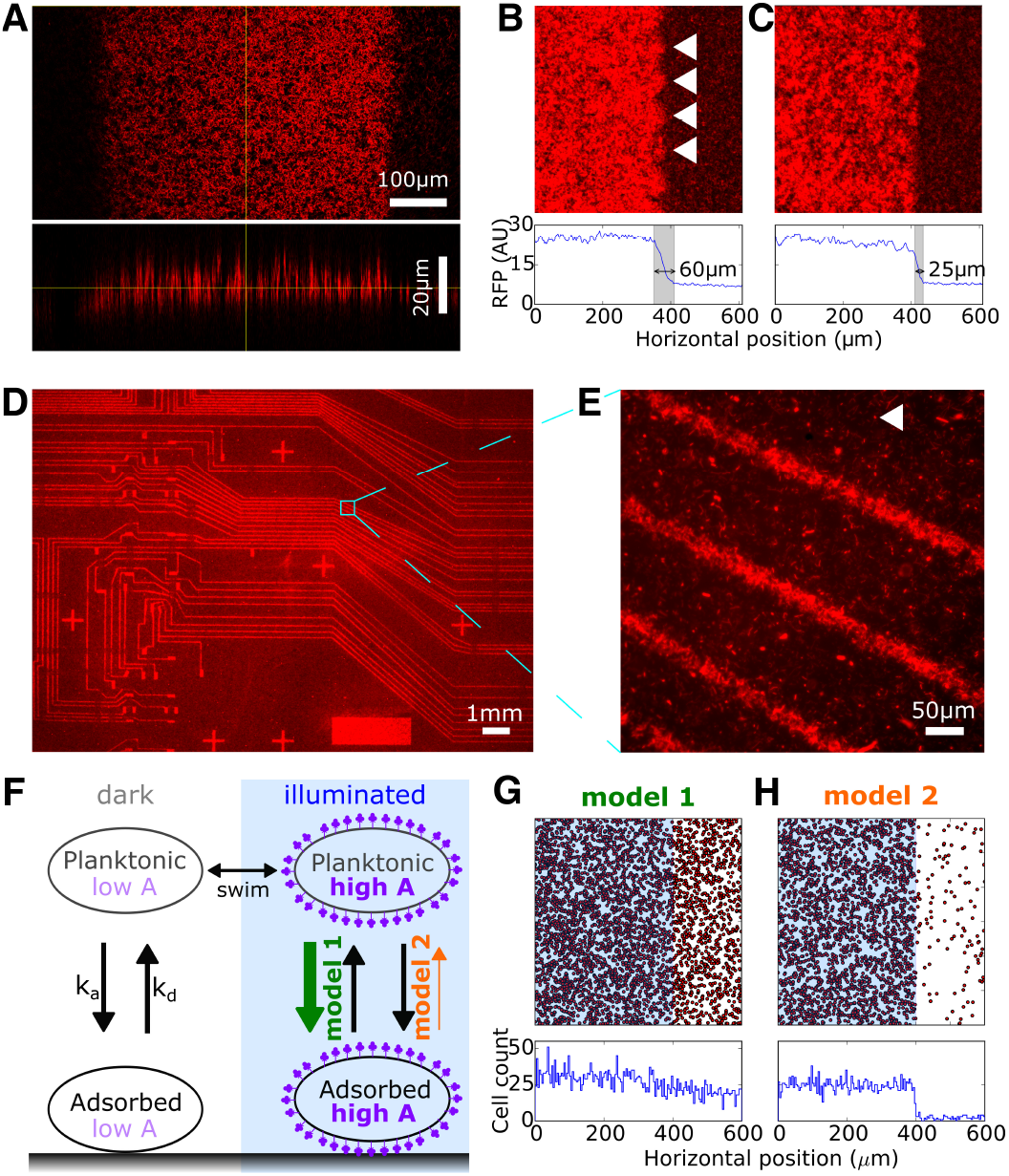
Biofilm Lithography enables patterning with a lateral resolution down to 25*μm*, which is explained by transient, light-independent cell-surface attachment followed by light activated adhesin expression. **(A)** Confocal microscopy reveals average biofilms thickness of 15*μm*, surface roughness coefficient of 0.33, and characteristic roughness length scale of 5.4*μm*. **(B)** Step-response analysis across light-dark boundary with high-resolution microscopy of striped illumination sample indicates ~ 60*μm* spatial resolution; white triangles point to artifacts due to projector pixels. **(C)** Using electrical tape as field-stop (as opposed to a projector) enables a spatial resolution of ~ 25*μm*. **(D)** Biofilm patterned over large area and at high resolution are possible with photomask (shown example was originally designed for patterning microfluidic channels). **(E)** Higher magnification imaging of **(D)** confirms feature sizes of ~ 25*μm*, white triangle points to example of a spurious individual cell. **(F)** Schematic of Monte-Carlo simulation with cell swimming, adsorption/desorption, and light-regulated levels of adhesin A. In **model 1**, increased adhesin increases adsorption rate of planktonic cells, whereas in **model 2**, increased adhesin decreases desorption rate of adsorbed cells increased adhesin. **(G)** No clear biofilm boundary can be observed at the light-dark border in model where optically regulated adhesin production increases adsorption rate *k_a_*. **(H)** Clear biofilm boundary can be observed in model where optically regulated adhesin production decreases desorption rate *k_d_*. (*G,H: Adsorbed cells are labelled red, illuminated region is marked in blue, histogram of cell positions plotted below*.)

The confocal images also reveal that the surface is not smooth, and we determine a surface roughness coefficient(33) of approximately 0.33 (Supplementary Information S5). This is in general agreement with the value of 0.31 estimated by assuming biofilm deposition to be a purely Poisson process (Supplementary Information S5). Using autocorrelation analysis, we derive an approximate length-scale for the surface roughness on the order of 5.7*μm* (Supplementary Information S5). We speculate that clustering on this length scale may be a result of cell division and intercellular Ag43 homodimerization leading to a different effective affinity for cell-cell vs. cell-surface binding.

Next, we determined the spatial resolution (smallest feature size) that could be patterned with this method by measuring the step-response of the red fluorescence signal across a light-dark illumination boundary. Using the striped illumination sample (Fig. 2a top), we estimate resolution of approximately 60*μm* (Fig. 3b). This approaches the optical resolution limit of the used projector setup with a pixel-pixel distance of approximately 80*μm*, which is also visible in corresponding periodic structural artifacts at this length scale (Fig. 3b, white triangles). To improve this resolution, we applied electrical tape to the bottom of the well-plate to act as a field-stop, and repeated the step-response analysis across the tape boundary. Based on this analysis, we estimate the spatial resolution to be 25*μm*, measured as the width of the transition region across the light-dark boundary (Fig. 3c).

To validate this point further, we taped a film photomask (originally designed for microfluidic circuit fabrication(34)) to the bottom of the culture chamber. This structure was faithfully recapitulated over the area of multiple mm^2^ (Fig. 3d) with feature sizes at the scale of 25*μm* (Fig. 3e). We noted that the feature sizes of the final patterned biofilm are reduced compared to the underlying photomask – i.e., 25*μm* wide stripes were observed in the biofilm corresponding to 75*μm* transparent stripes in the photomask (Supplementary Information S6). This reduction is roughly consistent with the 25*μm* light-dark transition region measured earlier, and suggests that the transition region lies on the illuminated side of the light-dark boundary. We also note spurious cells attached to the surface (Fig. 3e, white triangle), which along with the transition region and analysis on surface roughness lengthscales constitute a spatial resolution limit for ‘Biofilm Lithography’ as currently implemented.

### Biophysical modelling of biofilm formation explains high spatial resolution

Finally, we sought to understand mechanistically how MG1655/pDawn-Ag43 biofilm patterning can achieve high spatial resolution across illumination boundaries, given the inherent motility of the MG1655 strain. Our initial working hypothesis (**model 1**, Fig. 3f) was that pDawn-Ag43 works by optically stimulating Ag43 expression in freely swimming planktonic cells. The expression of surface appendages on cell membranes can help overcome cell-substrate electrostatic repulsion(35–37), which should lead to increased adsorption rate and higher levels of biofilm formation in illuminated regions. However, a back-of-the-envelope calculation suggests that this model is inconsistent with the observed 25*μm* spatial resolution. Motile *E. coli* have an effective diffusivity due to motility of *D_eff_* ~ 200*μm*^2^/s(38), while even conservative estimates of pDawn-Ag43 stimulation delay based only on gene expression time cannot fall below *T_delay_* ~ 100*s*(39–41). Bacterial swimming during the time delay between stimulation and attachment would blur features below a length scale of 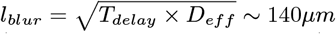, suggesting **model 1** is inconsistent with the observed spatial resolution given cell motility (Supplementary Information S7).

For another hypothesis (**model 2**, Fig. 3f), we note that pDawn-Ag43 mediated patterning occurs within the context of other biofilm-related processes – in particular, bacteria at a liquid-solid interface naturally exhibit reversible adsorption and desorption(42), switching between planktonic and transiently adsorbed subpopulations. Instead of only stimulating the planktonic subpopulation (by increasing their adsorption, as discussed above), pDawn-Ag43 could also have a significant effect on desorption rate of transiently adsorbed cells. Left unstimulated in the dark, these cells readily desorb from the surface(42), but in illuminated regions, membrane-expressed Ag43 reduces the desorption rate of adsorbed cells, effectively anchoring them to the surface(37). **Model 2** is consistent with high spatial resolution patterning that is not limited by bacterial motility since the adsorbed cells are immotile.

To quantitatively confirm these models, we developed a Monte-Carlo simulation of cell swimming and adsorption/desorption (Fig. 3f, see also Methods – Monte-Carlo Modeling). Individual cells are simulated to swim in a virtual culture chamber, one side of which is ‘illuminated’, causing increased production of adhesin ***A***. Bacterial cells adsorb and desorb from the surface with respective rates ***k_a_*** and ***k_d_***, which are a function of their adhesin expression level **A**. We simulated **model 1** by setting ***k_d_*** constant and ***k_a_*** an increasing function of ***A*** and observed no clear transition at the light-dark boundary (Fig. 3g). We simulated **model 2** by setting ***k_a_*** constant and ***k_d_*** a decreasing function of ***A*** and observed a clear increase in adsorbed cells in the illuminated region, with a sharp transition across the light-dark boundary (Fig. 3h). Notably, this model did not need to incorporate additional features such as increased adsorption, 3D cell-cell interactions, or a permanently attached cell state. Therefore, we establish that a biophysical model with just two key features, (1) optically controlled adhesin expression and (2) adhesin decreasing desorption rate of adsorbed cells, is sufficient to explain light-regulated biofilm formation with high spatial resolution despite bacterial motility. Additionally, the transition region in the simulations across the light-dark boundary in the simulations is less than 1*μm*, (Supplementary Information S8) – smaller than the experimentally observed 25*μm* transition region. This points to unaccounted-for experimental sources of noise such as colony growth/clustering, optical scattering, and 3D effects (e.g., cell stacking), on the other hand may inform future experimental strategies to improve spatial resolution even further.

## Discussion

In conclusion, we have developed a method (‘Biofilm Lithography’) that uses light-regulated adhesin expression (pDawn-Ag43) to quantitatively control biofilm formation and patterning with high spatial resolution. Compared to existing cell deposition and patterning approaches such as inkjet printing(12, 13), microcontact printing(14, 15), microfluidics(16), PDMS stenciling(17), patterned substrate modification(18–20), this method can pattern on a variety of surfaces, without the need for surface pre-patterning or pre-treatment, within enclosed chambers, over large areas, and at high spatial resolution. Rapid prototyping of different biofilm geometries is possible with low-cost digital projectors at resolutions of 60*μm*; resolutions down to 25*μm* can be reached with photomasks and likely also with more advanced optical projector setups (43, 44). This resolution represents an important step towards the engineering of biofilm communities, as natural biofilm microcolonies exist around this length scale(10). Our biophysical model suggests that the pDawn-Ag43 patterning mechanism works alongside natural surface adsorption/desorption in bacteria, and involves the stimulation of transiently adsorbed cells on the biofilm substrate towards a more permanently attached state. This insight gives additional support to the proposed role of Ag43 as an adhesin involved in biofilm maturation as opposed to initial surface adsorption(45). Ultimately, optogenetic patterning tools such as pDawn-Ag43 can be applied towards an improved understanding of naturally existing biofilms(46, 47), the design of synthetic microbial consortia(8), and new types of integrated diagnostic and microfluidic devices(48), with impact and trajectory that may potentially parallel that of silicon photolithography in the semiconductor industry(49, 50).

## Materials and Methods

### Plasmids and Bacterial Strains

MG1655 was obtained from the Coli Genetic Stock Center (CGSC strain #6300). pDawn-Ag43 was constructed using standard cloning techniques using pDawn (a gift from Andreas Moeglich, Addgene Plasmid #43796) plasmid as a starting point. First, Gibson Assembly was used to swap out the kanamycin marker for a less commonly used spectinomycin resistance marker. BamHI/XhoI restriction digest was then used to linearize the resulting plasmid at the multiple-cloning-site downstream of the λ promoter to create the backbone for pDawn-Ag43.

The coding sequence for Ag43 was obtained from the Biobricks IGEM distribution, part number *BBa_K*346007. We noted during construction that several PstI restriction enzyme sites remained in the coding sequence, making the part incompatible with the BioBrick Standard. Using a combination of site directed mutagenesis and DNA synthesis (IDT gBlock synthesis), these PstI sites were removed and replaced with silent mutations so as to not alter the final amino acid sequence. Using this now Biobrick-compatible part, we used standard Biobrick prefix/suffix assembly to add a medium strength ribosomal binding site (*BBa_B*0031 rbs) upstream of the Ag43 coding sequence. We then used BamHI/XhoI digestion/ligation to insert rbs-Ag43 into the backbone prepared earlier to create pDawn-Ag43.

A similar protocol was used to create pDawn-sfGFP, with a superfolderGFP coding sequence used in place of the Ag43 coding sequence. pBAD-Ag43 was created by using standard Biobrick prefix/suffix assembly to insert the Ag43 coding sequence downstream of an araC – pBAD expression vector (BioBricks part *BBa_I*0500). Sequences for pDawn-sfGFP, pDawn-Ag43, pBAD-Ag43 are available in the Supplementary Information online.

The plasmid for red fluorescent protein expression was obtained from the Biobricks iGEM distribution, part number *BBa_J*04450 – *pSB3T*5 backbone variant. This plasmid expresses mRFP from the Lac promoter using a strong ribosomal binding site, and also carries a tetracycline resistance marker on a low copy p15a origin of replication.

### Biofilm formation and patterning

E. coli strains were cultured to late log phase in LB broth under dark conditions (OD600 1.4, approximately 6h with shaking at 37°C after 1:1000 dilution of overnight culture). Media was supplemented with antibiotics as appropriate(50μg/mL for kanamycin and spectinomycin, 10μg/mL for tetracycline, and 100μg/mL for ampicillin). These cultures were then seeded onto non-tissue-culture treated polystyrene well plates at 1:100 dilution into M63 media supplemented with 0.2%w/v glucose and 0.1%w/v casamino acids.

For assays characterizing biofilm formation, 96-well blackwalled plates (Corning Costar 3631) were used. Patterning assays were performed in 6 or 12 well plates (Corning Falcon 351146/351143).

Well plates containing the biofilm cultures were taped to the ceiling of a 37°C incubator, ensuring the bottom surface of the well plates remained uncovered. An Ivation Pro4 Wireless Pocket Projector (IVPJPRO4) was secured below the ceiling of the incubator, pointing upwards toward the well plate on the incubator ceiling. The projector was connected via HDMI cable to a laptop through the incubator’s side access port, and Microsoft PowerPoint software was used to project blue light patterns.

Global illumination intensity was tuned by placing an adjustable neutral density filter (K&F concept AMSKU0124) at the aperture of the projector. Local illumination intensity was further tuned by taping thin neutral density filters from the Lee’s Filters Designer’s Edition Swatchbook (Lee’s Filters part SWB) to the bottom surface of the well-plates over specific wells to subject bacterial samples to a wide range of illumination intensities. Intensity of the projected pattern was measured using a Newport optical power meter with UV-vis photodetector (Newport 840C / 818-UV). Illumination time was adjusted on the software end through Microsoft PowerPoint.

For biofilm characterization experiments, biofilm cultures were placed in the incubator overnight (16h). Media was subsequently aspirated, and wells were gently washed twice with PBS. Wells were then stained with 0.1% crystal violet (Acros Organics 212120250) for 10 minutes, before another 2x wash with PBS. Wells were then allowed to dry before imaging, followed by A550nm quantification using 30% acetic acid solubilization as previously detailed(30).

To prepare samples for fluorescence microscopy, biofilm cultures were prepared using bacteria co-transformed with the *BBa_J*04450 – *pSB3T5* plasmid for mRFP expression. To pattern optical illumination for biofilm patterning experiments, various patterns were generated on the software end as PowerPoint illustrations. Cultures were incubated with patterned illumination as described above and rinsed twice with PBS before imaging under a wide-field fluorescence microscope. For the long-term culture experiments in PBS, cultures were prepared and patterned overnight, before media was aspirated, followed by PBS rinse. The sample was then imaged under a wide-field fluorescence microscope, before being left in PBS under ambient dark conditions for 3 days with daily imaging. The same protocpl was followed for long culture experiments in M63 media, except cell were maintained in M63 media with daily PBS rinse steps.

### Confocal Microscopy

To prepare cultures for confocal microscopy, a drop of self-hardening Shandon immunomount (Thermo Scientific 9990402) was dropped onto the cultured sample, before being covered with a glass coverslip (FisherScientific 12-545-81). The sample was then allowed to harden overnight at room temperature in the dark. The following day, the samples were imaged through the glass coverslip using a Leica Upright confocal microscope (Leica DMRXE), using a 20×/0.50 water immersion objective with an excitation line at 543 nm for mRFP.

### Monte-Carlo Modeling

The Monte-Carlo bacterial adhesion simulation was implemented in MATLAB using a forward Euler numerical approach, time discretized in *dt* = 100ms timesteps (repeats run with *dt* = *50ms* timesteps produced qualitatively identical results). Simulations were run for 16h over a (600*μm*)^2^ area. Within this area, a 400*μm* wide stripe on the left is ‘illuminated’ with blue light. Bacterial cells were initialized with a random position and velocity direction, with velocity magnitude in the range *υ* ≈ 14 ± 3*μm*/s(38). Cells were also initialized with a basal adhesin level of *A* = 1. Adhesin level is regulated by an ordinary differential equation.

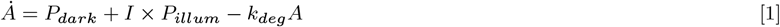

where I is a boolean variable representing whether the cell is being illuminated or not (in turn calculated based on the cell’s position). *P_dark_* and *P_illum_* represent the basal and light activated production rates of adhesion proteins respectively. These proteins are in turn degraded at a rate *k_deg_*. The parameter values *P_dark_* = 10^−3^s^−^, *P_illum_* = 10^−1^s^−1^, and *k_deg_* = 10^−3^s^−1^ were used so that in the dark cells revert to their original basal adhesin level *A* = 1 with characteristic protein turnover time on the order of tens of minutes(51), and in the light they increase their expression by two orders of magnitude (from *A_min_* = 1 to *A_max_* = 100) – approximately the reported dynamic range of pDawn(26). This ODE was numerically solved within the same forward-Euler loop as the overall simulation.

Simultaneously during each simulation time step, cell positions were updated at =based on velocity, and cells tumbled with a probability of *P_tumble_* = *f_tumble_* × *dt* based on a tumbling frequency of *f_tumble_* ≈ 1s^−1^(38). During a tumbling event, a cell’s velocity vector was reoriented randomly. Also at each time step, planktonic cells were adsorbed to the surface with a probability *P_adsorb_* = *k_a_* × *dt*, while adsorbed cells were desorbed with a probability *P_desorb_* = *k_d_* × *dt*. The rates of adsorption *k_a_* and desorption *k_d_* were dictated by adhesin level *A*.

In the first model, where cell adsorption is increased upon illumination, the relationships were set as *k_a_* = 10^−5^s^−1^ × *A* and *k_d_* = 10^−3^s^−1^. In the second model, where cell desorption was decreased upon illumination, the relationships were set as *k_a_* = 10^−5^s^−1^ and *k_d_* = 10^−3^s^−1^/*A*. The basal values for adsorption and desorption rate, *k_a_* = 10^−5^s^−1^ and *k_d_* = 10^−3^s^−1^, were derived from quantitative bacterial adsorption/desorption time measurements(42), and approximately correspond to planktonic cells adsorbing every few hours to the surface, after which they natively remain adsorbed for a few minutes if not stimulated, or a few hours if stimulated. Simulations were run for 576000 time steps, representing 16 hours of real time. The final state of the simulation was plotted with red dots marking the end position of the attached cells.

## ACKNOWLEDGMENTS

The authors thank D. Glass, A. Spormann, D. Endy, M. Covert, N. Cira, A. Choksi, S. Rajan, and A. Dvorak for helpful suggestions, the Spormann lab for access to their confocal microscope, and furthermore acknowledge the support from Stanford Bio-X Bowes and NSERC PGS fellowships, and the American Cancer Society (RSG-14-177-01). Author contributions: XJ and IHRK jointly conceived the project and wrote the paper, XJ performed experiments and analysis.

